# Spatial-extent inference for testing variance components in reliability and heritability studies

**DOI:** 10.1101/2023.04.19.537270

**Authors:** Ruyi Pan, Erin W. Dickie, Colin Hawco, Nancy Reid, Aristotle N. Voineskos, Jun Young Park

## Abstract

Clusterwise inference is a popular approach in neuroimaging to increase sensitivity, but most existing methods are currently restricted to the General Linear Model (GLM) for testing mean parameters. Statistical methods for testing *variance components*, which are critical in neuroimaging studies that involve estimation of narrow-sense heritability or test-retest reliability, are underdeveloped due to methodological and computational challenges, which would potentially lead to low power. We propose a fast and powerful test for variance components called CLEAN-V (**CLEAN** for testing **V**ariance components). CLEAN-V models the global spatial dependence structure of imaging data and computes a locally powerful variance component test statistic by data-adaptively pooling neighborhood information. Correction for multiple comparisons is achieved by permutations to control family-wise error rate (FWER). Through analysis of task-fMRI data from the Human Connectome Project across five tasks and comprehensive data-driven simulations, we show that CLEAN-V outperforms existing methods in detecting test-retest reliability and narrow-sense heritability with significantly improved power, with the detected areas aligning with activation maps. The computational efficiency of CLEAN-V also speaks of its practical utility, and it is available as an R package.

## 1. Introduction

Imaging biomarkers obtained by magnetic resonance imaging (MRI) are critical to the understanding of brain functions and structures and their relationship to the diagnosis of brain disorders. Among these, functional MRI (fMRI) has been increasingly used to measure brain activation in response to experimental tasks to understand human cognition and behavior. However, recent studies using large neuroimaging databases (e.g., the Human Connectome Project (HCP) and Adolescent Brain Cognitive Development (ABCD) study) have shown poor test-retest reliability of task-fMRI measures in most brain regions [1, 2]. Since test-retest reliability provides a measure of consistency and stability of data obtained repeatedly over time, the lack of reliability of task-fMRI measures has raised concerns about current efforts using fMRI data to find brain-behavior associations [3]. Blokland et al. [4], with 319 twins in their study, discovered several regions of the brain highlighting heritable fMRI activations, but only with a loose cluster defining threshold (uncorrected voxel-wise *p <* 0.05), which might be prone to inflate false positives. These recent findings in neuroimaging motivate a need for a powerful statistical method for neuroimaging studies that involve test-retest reliability or heritability.

Conducting inference for test-retest reliability and heritability is fundamentally a variance component problem in statistics [5] because both measures can be expressed in terms of random effects for modeling dependencies between observed data (images). However, most existing variance component methods in neuroimaging literature estimate and test each variance component separately (e.g., massive univariate analysis). In contrast to testing for mean parameters, testing for variance component parameters is more challenging due to model specification and intensive computation to address spatial dependence structures. Recently, Risk and Zhu [6] showed that spatial modeling of variance components would lead to improved heritability estimation. However, their method also suffers from computational costs, and conducting statistical inference with their method is not straightforward, as the authors also noted.

We hypothesize that leveraging spatial dependencies in testing for the test-retest reliability or heritability would increase the replicability of identified regions [7]. The spatial-extent inference has shown its utility to increase sensitivity in neuroimaging compared to simple massive univariate analysis. In the context of the General Linear Model (GLM), clusterwise inference improves sensitivity through the convolution of univariate test statistics in the spatial domain, which is exemplified by the threshold-free clusterwise enhancement (TFCE) [8]. Park et al. [9] showed that combining the test statistics across neighboring vertices yielded high statistical power in linear mixed models as well. Also, spatial Bayesian GLMs that parametrically model spatial dependencies of the mean parameters resulted in more accurate and reliable estimates [10, 11]. Bernal-Rusiel et al. [12] and Risk et al. [13] showed that modeling of the local spatial dependence of the residual noise term using a Gaussian process results in increased statistical power. More recently, the CLEAN method proposed by Park and Fiecas [14] showed that additional gain in sensitivity is achieved in GLMs by cluster-enhanced statistic, which is constructed from multivariate test statistics by applying spatial Gaussian process to the residual noise. This idea has been extended to CLEAN-R [15] in testing and localizing correspondence between two imaging modalities by modeling modality-specific spatial autocorrelation and constructing cluster-enhanced test statistics for correlation parameters. CLEAN and CLEAN-R highlight the importance of using two aspects of spatial dependencies to improve sensitivity.

The aim of this paper is to extend spatial-extent inference to testing variance components, which have been underdeveloped in neuroimaging due to their methodological and computational challenges. To overcome the limitations of massive univariate analysis in variance component testing, we propose a new powerful method called CLEAN-V (**CLEAN** for testing **V**ariance components) that tests and localizes variance components at the vertex level. CLEAN-V shares the same underlying mechanism as CLEAN [14] and CLEAN-R [15] in improving statistical power: (i) an explicit spatial autocorrelation modeling of neuroimaging data, (ii) effective spatial enhancement of test statistics, and (iii) a fast resampling procedure for making statistical inferences. However, CLEAN and CLEAN-R are primarily developed for testing GLM parameters or intermodal correlations, and their direct applications to test variance components in test-retest reliability and heritability studies are limited. Therefore, the proposed method aims to bridge this gap and provide a robust solution for testing and localizing variance components in neuroimaging studies.

The rest of the paper is organized as follows. In Section 2, we describe the details of CLEAN-V and compare it to the existing variance component testing methods. In Section 3, we compare the performance of CLEAN-V to other existing methods by conducting comprehensive data-driven simulation studies. We apply CLEAN-V to multiple tasks from Human Connectome Project (HCP) to compare statistically significant test-retest reliability and heritability regions across different experimental tasks. We conclude with discussion in Section 4.

## 2. Methods

### 2.1. Notation and model specifications

We consider a dataset consisting of subject-level continuous neuroimaging data. In this paper, we focus on activation levels in task-fMRI measured on the cortical surface, although other neuroimaging data types such as cortical thickness and cerebral blood flow that are mapped onto the cortex are also applicable. Task activation for each individual at each location is obtained by applying GLM to the observed blood oxygenation level-dependent (BOLD) signal on the convolution of the stimulus function with the hemodynamic response function [16, 17].

Consider *N* total images where each image has *V* vertices. Note that *N* does not necessarily denote subjects in reliability studies, where more than two scans from a subject are obtained. Let *i* = 1, …, *N* be the indices for images, and *v* = 1, …, *V* be the indices for vertices in a cortical surface. The observed imaging data **y**(*v*) = (*y*_1_(*v*), *y*_2_(*v*), …, *y*_*N*_ (*v*))^′^ are modelled by

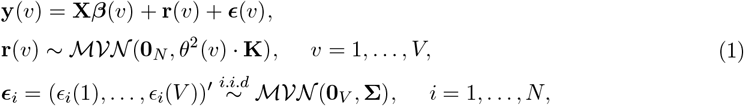

where **r**(*v*)= (*r*_1_(*v*), …, *r*_*N*_ (*v*))^′^ and ***ϵ***(*v*) = (*ϵ*_1_(*v*), …, *ϵ*_*N*_ (*v*))′ are assumed to be independent of each other. Please refer to Figure 1 for illustrations of the model specification. Here, **X** is an *N* × *p* nuisance covariates matrix (e.g., age): each row of the matrix (**x**′*i*) is a *p*-dimensional nuisance covariates vector for the *i*th image. Under this model, **r**(*v*) characterizes the dependence among all *N* images at vertex *v* (e.g., vertex-level test-retest reliability or heritability) through **K**. We note that the dependence of images at vertex *v* is captured through *θ*^2^(*v*). Without any true reliable (or heritable) features, we have *θ*^2^(*v*) = 0 for all *v*, and images are considered independent.

**Figure 1:**
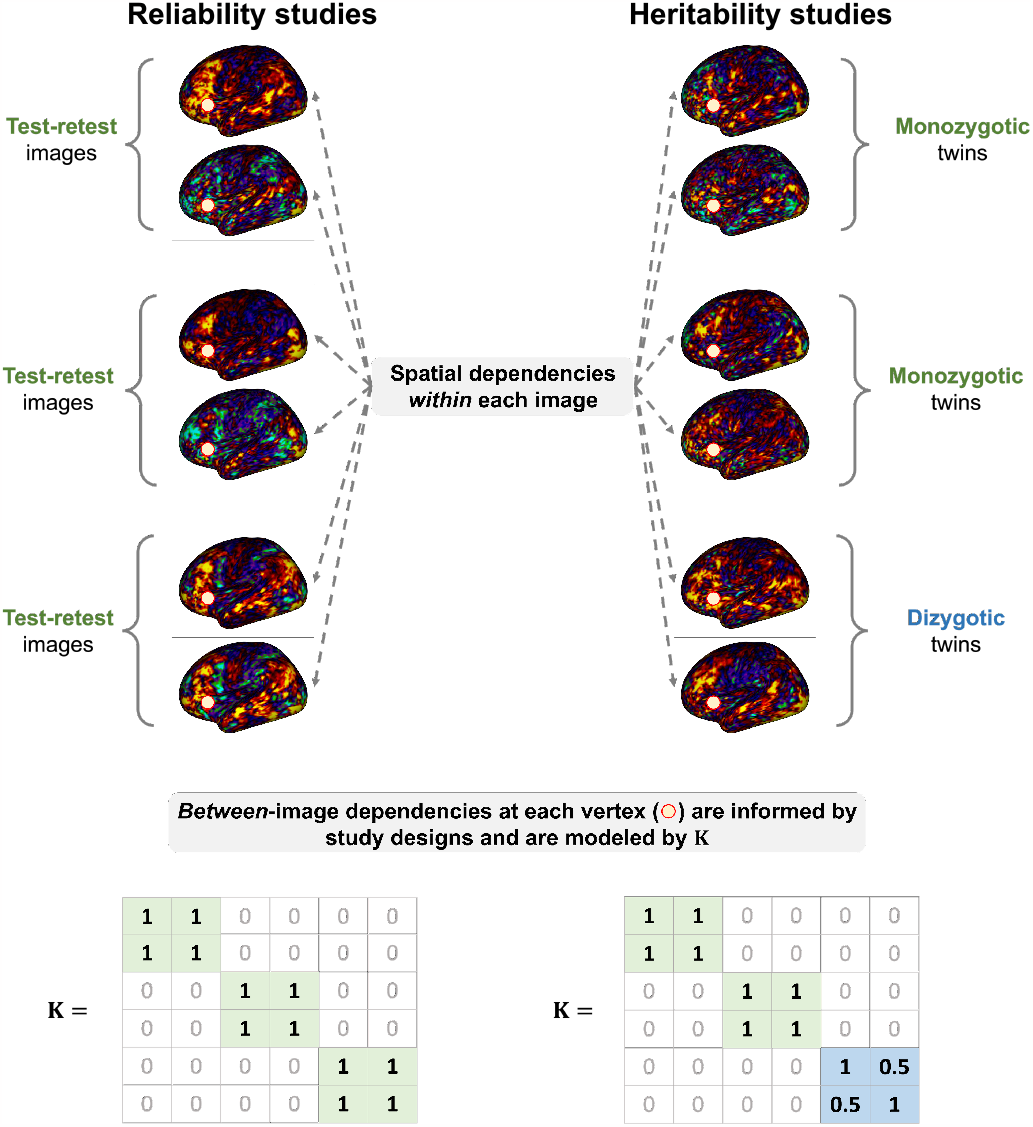
Visualization of two sources of dependencies in reliability and heritability studies. Our model specification captures the spatial dependencies within each image (top) as well as between-image dependencies at each vertex informed by study designs, which are modeled by **K**.

In this model, **Σ**, which is the covariance of the *i*th residual image ***ϵ***_*i*_ = (*ϵ*_*i*_(1), …, *ϵ*_*i*_(*V*))′, characterizes both spatial and non-spatial variations within images. We use **b**_*i*_ = (*b*_*i*_(1), …, *b*_*i*_(*V*))′ to characterize the variation due to spatial autocorrelation, whose covariance is modeled by *σ*^2^**Φ**(*ϕ*, 𝒟) with a scale parameter *σ*^2^ and a predefined spatial autocorrelation function (SACF) with a parameter *ϕ >* 0 and a *V* × *V* pairwise geodesic distance matrix 𝒟. We also use ***δ***_*i*_ = (*z*_*i*_(1), …, *z*_*i*_(*V*))′ to characterize the non-spatial variations, or white noise, with its covariance *τ* ^2^**I**_*V*_. The assumption that **b**_*i*_ and ***δ***_*i*_ are independent leads to the **Σ** = *σ*^2^**Φ**(*ϕ*, 𝒟) + *τ* ^2^**I**_*V*_, which follows ***ϵ***_*i*_ = **b**_*i*_ + ***δ***_*i*_ with

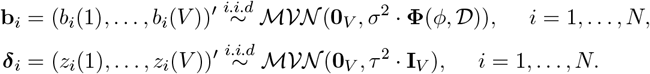

Following exploratory data analyses of Park and Fiecas [14] and Weinstein et al. [15], we assume the exponential spatial autocorrelation function (SACF) for modeling the contrast of parameter estimates in task-fMRI. Thus, the (*v, v*^∗^)th element of **Φ**(*ϕ*, 𝒟) can be expressed by exp(−*ϕ* · *d*_*v,v*∗_) where *d*_*v,v*∗_ is the geodesic distance between vertices *v* and *v*^∗^.

From the model specification, the null hypothesis of interest is

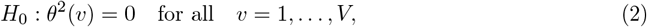

which implies that there is no between-image dependencies. The model shows flexibility in specifying reliability and heritability structures through the specification of the dependence structure **K**. In testing for test-retest reliability, we specify the (*i, i*^∗^)th element of **K** with 1 if images *i* and *i*^∗^ are from the same subject and 0 otherwise. In testing narrow-sense heritability in twin studies, the (*i, i*^∗^)th element of **K** with 1 if the subjects *i* and *i*^∗^ are monozygotic twins, 0.5 if they are dizygotic twins, 0 otherwise. In larger pedigree studies, **K** becomes a genetic relationship matrix.

### 2.2. Existing approaches

Existing approaches in testing for variance components are mostly massive univariate analyses where spatial dependencies of the imaging features are not considered. Specifically, the **b**(*v*) terms (spatial variations) are dropped, and the model reduces to a massive vertex-wise model as

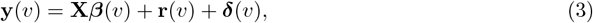

where the null hypothesis remains the same as (2). Note that, from this vertex-level model, narrow-sense heritability is defined by variance due to additive genetic effects divided by the total variance, and test-retest reliability is defined by intraclass correlation coefficient (ICC), both of which giving the same parametrization as *θ*^2^(*v*)*/*(*θ*^2^(*v*) + *τ* ^2^(*v*)) based on study designs.

Ganjgahi et al. [18] used the massive univariate method to carry out voxel-wise heritability inference and estimation with the likelihood ratio test (LRT), Wald test, and score test. Their study results showed that the most efficient method, the score test, produced more conservative inferences. They argued that it is likely due to untenable 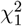mixture approximation even if all of the tests have a common asymptotic result following a 50:50 mixture of 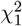 distribution and 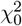 (i.e., point mass at 0).

Ge et al. [19] proposed massively expedited genome-wide heritability analysis (MEGHA) by fitting the model (3) to all vertices with brain structure measures. They used a scaled variance-component score test statistic. The variance-component score test statistic is also known as the sequence kernel association test (SKAT) statistic with a linear kernel in genome-wide association studies (GWASs) [20]. SKAT is a computationally efficient method to test the association between genetic variants since the computation of the test statistics only requires parameter estimates obtained under the null model. The statistic has the form

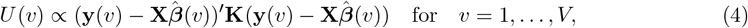

Where 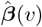is estimated under the null hypothesis (2). Wu et al. [20] showed that the SKAT statistic follows a mixture chi-square distribution under *H*_0_, where the weights of the mixture chi-squares are determined by the eigenvalues of **QKQ**′ and **Q** is a Cholesky decomposition of 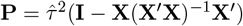 that satisfies **P** = **QQ**′. Wu et al. [20] also showed that it is the locally most powerful test compared to the log-likelihood ratio test.

### 2.3. Proposed method: CLEAN-V (‘CLEAN’ for testing ‘V’ariance components)

From the spatial model specified in Equation (1), the score-based variance component test statistic for vertex *v* is

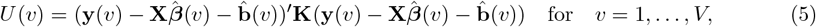

with 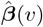and 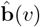estimated under *H*_0_. Importantly, fitting the null model without any *θ*^2^(*v*)s is equivalent to fitting CLEAN for GLM [14] because dependencies of images are modeled by **r**(*v*) only. Therefore, we first use ordinary least squares to estimate ***β***(*v*) for each *v* separately and use residual images 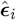 to obtain consistent estimates *σ*^2^, *ϕ* and *τ* ^2^ via covariance regression analysis [21, 14] by minimizing

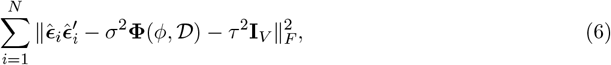

where 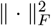 is the squared Frobenius norm of a matrix (Appendix A of Park and Fiecas [14]). Then, we use the parameter estimates to construct 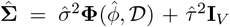 and obtain 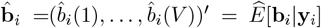 under the null model. It can also be shown that *U* (*v*) is also a score-based variance component test statistic under the working (spatial) covariance matrix, which implies that *U* (*v*) from Equation (5) also follows a mixture chi-square distribution (Supplementary material A).

We construct CLEAN-type cluster-enhanced test statistics in CLEAN-V for its powerfulness and easy implementation in the mesh surface. Compared to other types of cluster enhancement (e.g., TFCE [8]), the CLEAN-type cluster enhancement is constructed from aggregating test statistics from the local neighborhood [14]. To construct cluster-enhanced test statistics using *U* (*v*) in Equation (5), we first define *N*_*r*_(*v*) as the collection of all vertices within the radius *r* neighborhood of a central vertex *v* (i.e., {*v*^∗^ : *d*_*v,v*∗_ ≤ *r*}). Then, we use all the *U* (*v*^∗^) from vertices *v*^∗^ within *N*_*r*_(*v*) to construct a cluster-enhanced test statistic for the vertex *v*. For a radius *r*, we compute the standardized sum of the variance-component score test statistics within *N*_*r*_(*v*):

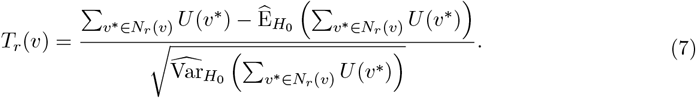

In practice, we obtain the null distribution of 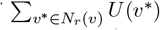 by permutation and use empirical permuted test statistics to obtain 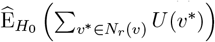 and 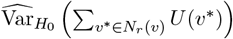 by the sample mean and sample variance (Section 2.4). Similar to how we used the quadratic form to show that *U* (*v*^∗^) follows a mixture of chi-square distribution, 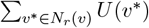 also follows a mixture of chi-squares distribution under the null hypothesis (Supplementary material A). A mixture chi-square distribution can be approximated by normal distribution according to the Lyapunov central limit theorem (Supplementary material A). Therefore, after standardization in (7), *T*_*r*_(*v*) approximately follows the standard normal distribution for each *r* and *v*. This transformation makes sure that each test statistic is mapped onto the same standard normal scale, which allows for data-adaptive testing based on the true areas of signals. Figure 2 is given for illustration.

**Figure 2:**
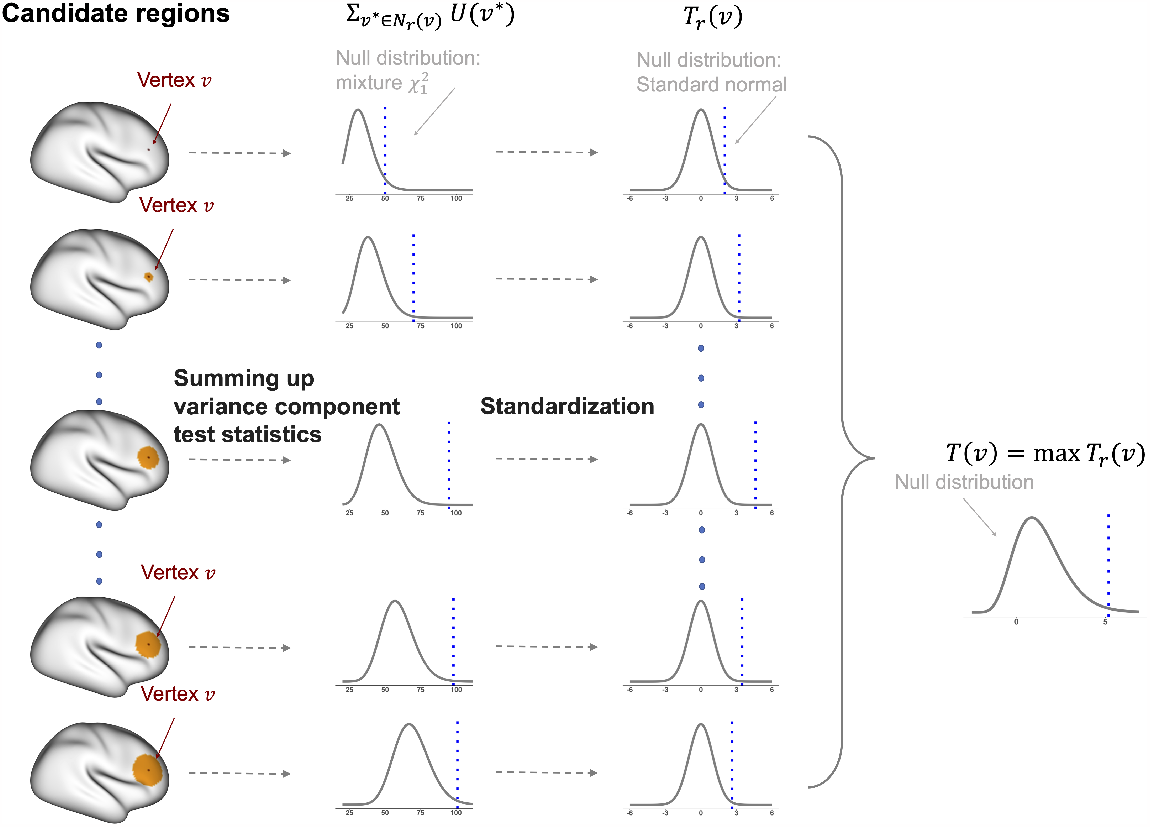
The process to construct clusterwise-enhanced statistics in CLEAN-V.

Statistically, the way CLEAN-V achieves high power aligns with CLEAN [14] developed for GLM. In CLEAN, *T*_*r*_(*v*) is defined by assuming that the regression coefficients of interest in *N*_*r*_(*v*) are the same [14]. Similarly, it can be shown that *T*_*r*_(*v*) in CLEAN-V is constructed by assuming that *θ*^2^(*v*^∗^)s in *N*_*r*_(*v*) are the same (Supplementary material B). Since the SKAT statistics are known to be locally most powerful, combining test statistics within a neighborhood strengthens the evidence against the null hypothesis when the strength of signals remains consistent within the neighbor. This property, along with Supplementary material B, makes the proposed CLEAN-type enhancement particularly powerful in CLEAN-V. Li et al. [22] used a similar approach, called Quadratic Scan (Q-SCAN) statistic, to identify the sizes and locations of signal regions in genomewide association studies.

In CLEAN-V, we build a map of test statistics by adopting an adaptive method to benefit from the ‘ideal’ local neighbor *N*_*r*_(*v*) for each *v*. Since the best size of the neighborhood area for each vertex might not be the same, we don’t fix the radius *r*. Instead, we consider different clusters by choosing different radii *rs* from *r*_1_ up to *r*max and choose the maximum *T*_*r*_(*v*) as the adaptive cluster-enhanced test statistic for vertex *v*:

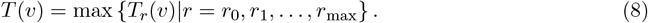

In this way, we can find the closely optimal cluster size for each vertex, which is useful as true areas of signals are unknown *a priori*. Following Park and Fiecas [14] and Weinstein et al. [15], we consider *r*_1_ = 1mm, *r*_2_ = 2mm, …, *r*_max_ = 20mm as a default in this paper, although *r*_max_ can be chosen flexibly based on research questions and contexts. The radius *r*_0_ = 0mm is included in the radii set since it guarantees that original vertex-level test statistics can also be considered in the maximization process.

### 2.4. FWER control with permutation

A critical value *t*_*α*_ is needed to control the family-wise error rate (FWER) at level *α*. We consider the test statistic for *H*_0_ as

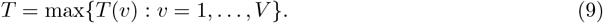

Permutation allows us to generate an empirical distribution of test statistics under the null hypothesis. In our setting, it can be achieved by shuffling 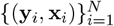 while keeping the dependence matrix **K** unchanged. Note that this is equivalent to shuffling 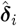 from the null model across images, which in practice does not require any refitting of the mean model and the covariance model (Equation (6)). For the *b*th permutation (*b* = 1, …, *B*), we construct the permuted variance-component test statistics *U* ^(*b*)^(*v*). Then, we use Equation (7) to construct the permuted single cluster-enhanced test statistic 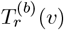. *T* ^(*b*)^(*v*) and *T* ^(*b*)^ follow Equations (8) and (9). Finally, we construct the empirical null distribution by the set {*T* ^(1)^, …, *T* ^(*B*)^} and select the (1 − *α*)th quantile as *t*_*α*_. Statistical significance is achieved at *α* × 100% FWER when *T > t*_*α*_.

### 2.5. Localizing areas of significance

The set of vertices *v* with statistically significant *θ*^2^(*v*) *>* 0 is identified by vertices whose *T* (*v*) exceeds *t*_*α*_.

### 2.6. Notes on the computational efficiency

Considering the complicated covariance structures implied in Equation (1), CLEAN-V reduces the computational cost dramatically because CLEAN-V uses score-based test statistics that require the estimation *σ*^2^, *τ* ^2^, *ϕ* only. Also, the permutation does not even require estimating *σ*^2^, *τ* ^2^, *ϕ* repeatedly but matrix multiplications from the null model only. We note that other methods, such as likelihood-ratio test or Wald test used by Ganjgahi et al. [18] would require the estimation of *θ*^2^(1), …, *θ*^2^(*V*), *σ*^2^, *τ* ^2^, *ϕ* in the original data and each permutation when extended to the spatial autocorrelation modeling, which would be computationally infeasible. In addition, in CLEAN-V, computational costs for obtaining *σ*^2^, *τ* ^2^, *ϕ* under the null model are relaxed by using a method-of-moment approach (Equation (6),[21]), and the calculation of 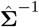 for a large *V* is relaxed by using the nearest neighbor Gaussian process approximation [23].

## 3. Data analysis and data-driven simulations

### 3.1. Data preparation

We collected task-fMRI data from the Human Connectome Project (1200 Subjects Data Release). We focus on 5 experimental tasks: relational processing, emotional processing, social cognition, language processing, and gambling decision-making. Specific contrasts we used for each task are summarized in Table 1. Barch et al. [24] provide a more detailed description of each task. There are more than 1000 subjects in total who completed the tasks and, among these, 44 subjects who completed tasks twice. Additionally, there are 171 twin pairs (*N* = 342 images), of which 68 pairs are dizygotic twins and 103 pairs are monozygotic twins. There are 80 and 116 female twins correspondingly. The average ages for dizygotic and monozygotic twins are 29.16 (SD: 3.42) and 29.29 (SD: 3.37) years, respectively. The task-fMRI data were processed through the HCP’s minimal surface processing pipeline with 2mm surface smoothing [25]. We used the R package ciftiTools [26] to sample 10242 vertices from approximately 32000 vertices resulting in 9394 and 9398 cortex vertices in the left and right hemispheres, respectively, after excluding 848 and 844 vertices in the medial wall in each hemisphere.

**Table 1:**
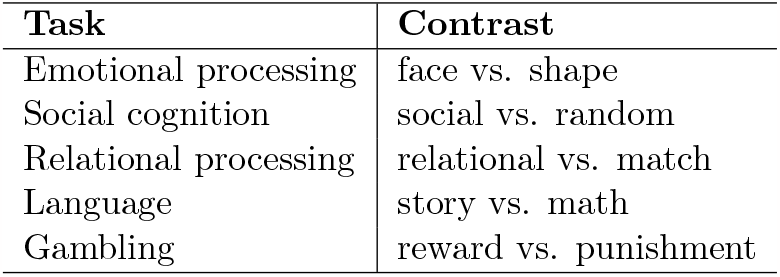
The list of selected contrasts for each task used in our analysis.

| Task | Contrast |
| --- | --- |
| Emotional processing | face vs. shape |
| Social cognition | social vs. random |
| Relational processing | relational vs. match |
| Language | story vs. math |
| Gambling | reward vs. punishment |

### 3.2. Competitors

In our data analysis and simulations, we compared CLEAN-V to three competitors, all of which are considered as simplified versions of CLEAN-V. Specifically,

1. **CLEAN-V without spatial correlation**: It constructs cluster-enhanced test statistics *T* (*v*) based on the model without considering spatial autocorrelations (Equation (3)). For a fair comparison, we used the same neighbor set as CLEAN-V to construct cluster-enhanced test statistics.
2. **CLEAN-V without cluster enhancement**: It models spatial autocorrelation of the noise following Equation (1), but it does not construct cluster-enhanced statistic. It is equivalent to setting *r* = 0 only for *T*_*r*_(*v*) (i.e., the vertex itself).
3. **Massive-univariate analysis**: This method neither models spatial autocorrelation nor constructs cluster-enhanced test statistics, which is equivalent to MEGHA [19].

All these competitors used the same permutation step outlined in Section 2.4. We included *CLEANV without spatial correation* and *CLEAN-V without cluster enhancement* to examine the effect of modeling the spatial structure of features and the impact of the novel clusterwise enhancement method, separately.

### 3.3. Data analysis

#### 3.3.1. Test-retest reliability and heritability for five different cognition tasks

We first fitted CLEAN-V to (i) 44 subjects with test-retest images to obtain the test-retest reliability map and (ii) 171 twin pairs to obtain the heritability map, with FWER controlled at 5%. These maps are shown in Figure 3(a). CLEAN-V detected wide areas of statistically significant test-retest reliability on the brain surface for most of the task-fMRI. However, we obtained different areas of significance for different tasks. The language task showed strong evidence of test-retest reliability, but the gambling task showed tiny and sparse test-retest reliability. However, the five tasks displayed a common result: anterior central gyrus and postcentral gyrus showed low or no test-retest reliability.

**Figure 3:**
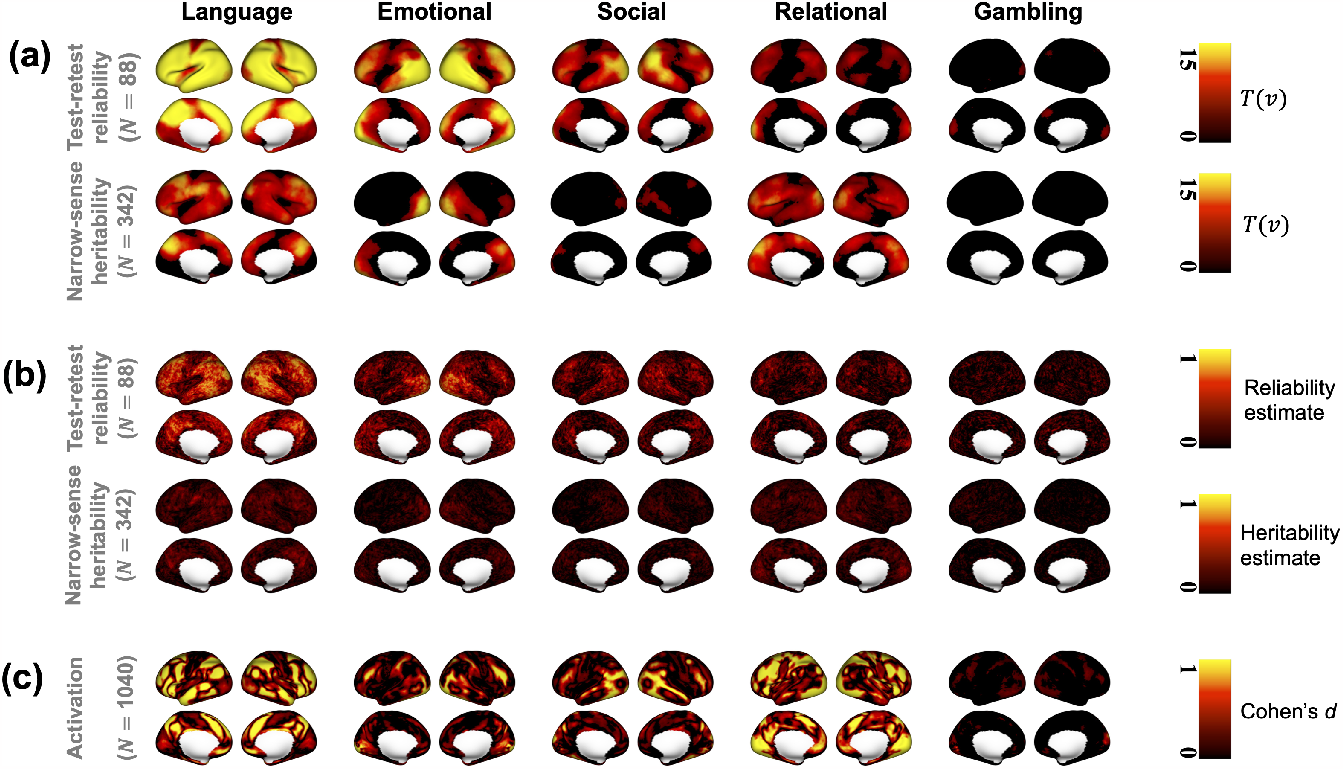
(a) CLEAN-V’s localization results (*T* (*v*)s) for the five cognition tasks. For visualizations, *T* (*v*)s were shrunk to zero if they were less than the threshold *t*_0.05_. (b) The effect size estimates of the test-retest reliability and narrow-sense heritability 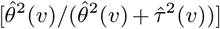, estimated based on the vertex-level (non-spatial) model in Equation (3). (c) The effect size (Cohen’s *d*) maps for the group-level activation, which is thresholded at 0.2 for visualizations.

We also provided estimates of test-retest reliability and narrow-sense heritability (Figure 3(b)) for comparison. Their effect size computed by the massive univariate method was small and specific to tasks, as expected. However, CLEAN-V detected test-retest reliability and narrow-sense heritability with high significance even if the estimates were small, and the detected regions corresponded well to the effect size estimates. Also, By qualitatively comparing the test-retest reliability maps to the activation maps (Cohen’s *d*; Figure 3(c)), we found that there is a strong correspondence between the strength of reliability and activation [27]. Specifically, the language tasks showed a strong positive and negative activation map which agrees with our results: the strongest evidence of test-retest reliability came from the language task. The weak evidence of activations in gambling tasks also agrees with the test-retest reliability map of CLEAN-V.

#### 3.3.2. Localization results from CLEAN-V and its competitors

Figure 4 shows the CLEAN-V and the competitors’ binarized localization results of both testretest reliability and narrow-sense heritability for the emotional processing task. CLEAN-V without cluster enhancement produced almost the same result as the massive univariate analysis, where very limited number of vertices were detected. Compared to these two methods, CLEAN-V without spatial correlation detected obviously more regions, which shows using cluster-enhanced test statistics improved signal detection. CLEAN-V detected most of the areas for the reliability and heritability, which validates the benefits of cluster-enhanced tests statistics and spatial autocorrelation modeling. The analysis results for other tasks are provided in Figures S1 and S2 in the supplementary material C, and the performances were analogous to the emotional task which supported the most sensitive performance of CLEAN-V and the least sensitive performance of massive univariate analysis.

**Figure 4:**
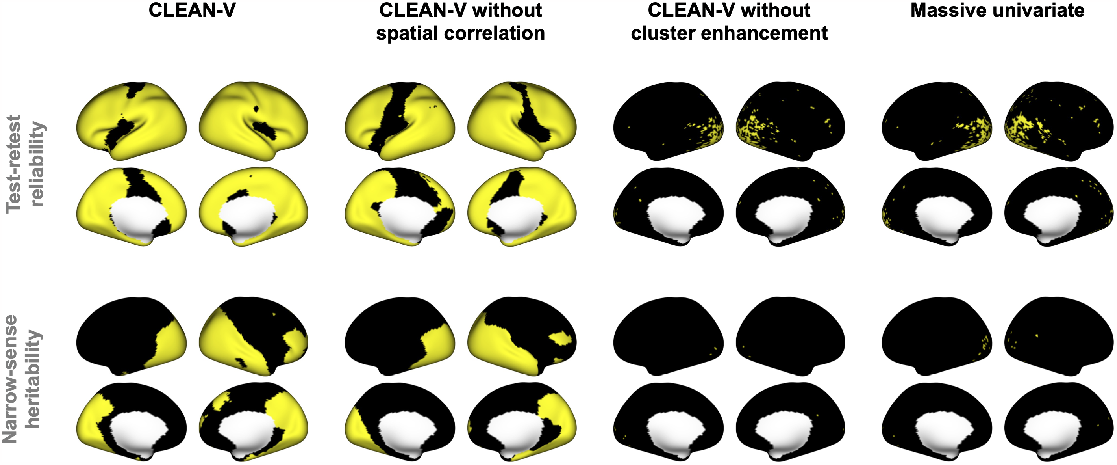
Localization results from CLEAN-V and its competitors for emotional task are displayed on the inflated surface. The yellow vertices denote for statistically significant vertices detected by each method.

#### 3.3.3. Comparisons to massive univariate analysis with smoothed images

In this section, we also conducted massive univariate analyses (MEGHA) with different levels of surface smoothing (e.g., 5mm, 8mm, 10mm). We compared them to CLEAN-V applied to the original data but with *r*_max_ = 0mm, 5mm, 8mm, 10mm, although we acknowledge that a direct comparison between ‘smoothing radius’ in massive univariate analysis and *r*_max_ in CLEAN-V is not feasible. From Figure 5, massive univariate analysis applied to presmoothed data improved the number of detected vertices. For CLEAN-V, it also consistently increased the detected regions as *r*_max_ increased. At the same time, it suggests that even a small *r*_max_ (e.g., 5mm) seems to outperform massive univariate analyses with different smoothing levels. These results demonstrate the relative efficiency of CLEAN-V.

**Figure 5:**
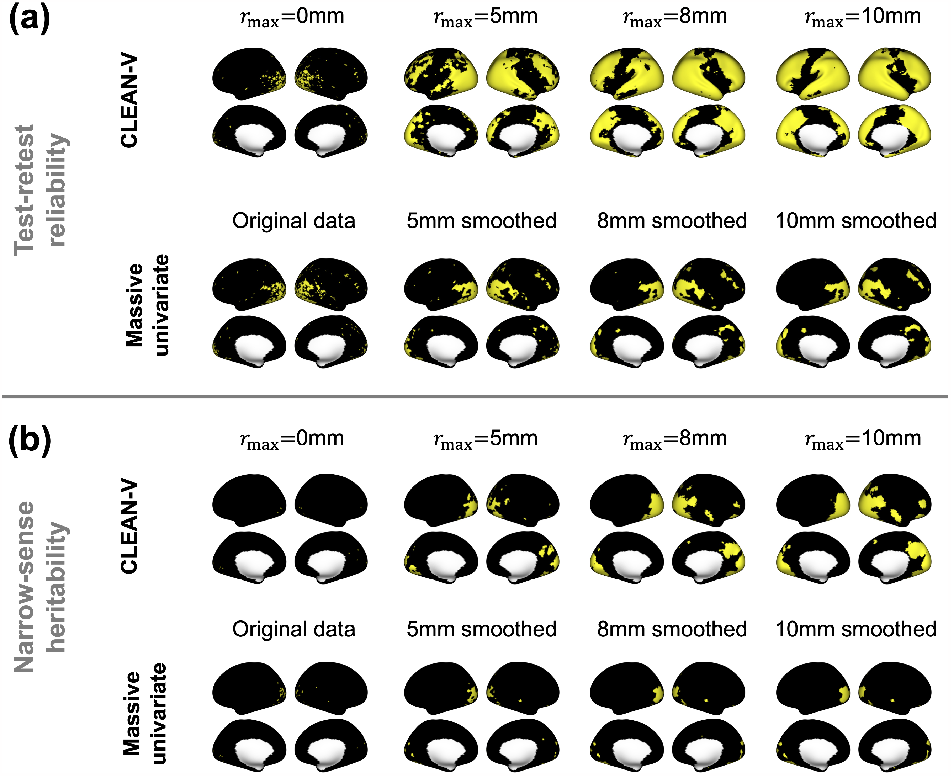
Binarized localization results from CLEAN-V and massive univariate analysis for emotional task. CLEAN-V models the original data but with different maximum radius *r*_max_ for cluster-wise enhancement process. MEGHA models smoothed data under different levels of surface smoothing.

A cautionary note is warranted. Both imaging smoothing and cluster enhancement involve a tradeoff between sensitivity and specificity. Therefore, just as choosing the optimal smoothing level is an open question, choosing an optimal *r*_max_ in CLEAN-V, similar to CLEAN and CLEAN-R, is an open question which should be determined carefully based on expected signal-to-noise ratios, sample sizes, and research purposes.

### 3.4. Data-driven simulations

#### 3.4.1. Setup

Motivated by existing efforts in neuroimaging literature [14, 15, 28, 29], we used task-fMRI data from HCP to design data-driven simulations to evaluate the performance of CLEAN-V compared to the competitors. It aligns with the approach of Eklund et al. [30], which used real data (restingstate fMRI) to evaluate false positive rates of different statistical methods for task-fMRI. Also, this approach is particularly useful in evaluating the robustness of CLEAN-V under potential model misspecifications (in covariance structures) because it assumes stationarity and isotropy of underlying spatial autocorrelation with the exponential SACF, which may not hold. For our datadriven simulations, we used images from the emotional task to construct ‘null’ data and ‘signal’ data with different signal-to-noise ratios (SNRs), which are described below.

#### 3.4.2. Simulating null data

The null data consists of independent images only. Among all subjects in HCP, we only considered test images from non-twin subjects to construct ‘pseudo’ test-retest pairs or twin pairs. For example, a set of test-retest images for a subject was generated by randomly sampling two images from test images from different independent (non-twin) subjects. Since there are no true test-retest pairs or twins in the null data, we expect no test-retest reliability or narrow-sense heritability.

#### 3.4.3. Simulating test-retest data with different signal-to-noise ratios

We fixed the number of subjects to be 20 (*N* = 40 images) throughout the simulation studies for test-retest reliability, and we changed the proportion of subjects whose test and retest images are sampled (without replacement) from the 44 subjects available from HCP. The remaining subjects’ images were sampled from test images from different subjects from HCP. We expect stronger SNR when more test-retest pairs are included. These evaluations are shown in Section 3.5.1.

#### 3.4.4. Simulating twins data with different signal-to-noise ratios

The simulation process for twins data is the same as the procedure for test-retest data except for the larger sample size. We fixed the number of twin pairs to be 60 (*N* = 120 images) as the overall effect size of narrow-sense heritability is smaller than test-retest reliability. We changed the proportion of true twins whose images were sampled from the 171 twins. The rest of the image pairs were sampled from different subjects. These evaluations are also shown in Section 3.5.1.

### 3.5. Data-driven simulation results

In each scenario, we generated 1000 sets of simulated data to evaluate performances.

#### 3.5.1. Power analysis and family-wise error rate

Figure 6(a) shows the empirical FWER of the CLEAN-V, which demonstrates that CLEAN-V and its competitors controlled FWER accurately at 0.05 under different maximum radii *r*_max_ of cluster enhancement for testing both test-retest reliability and narrow-sense heritability.

**Figure 6:**
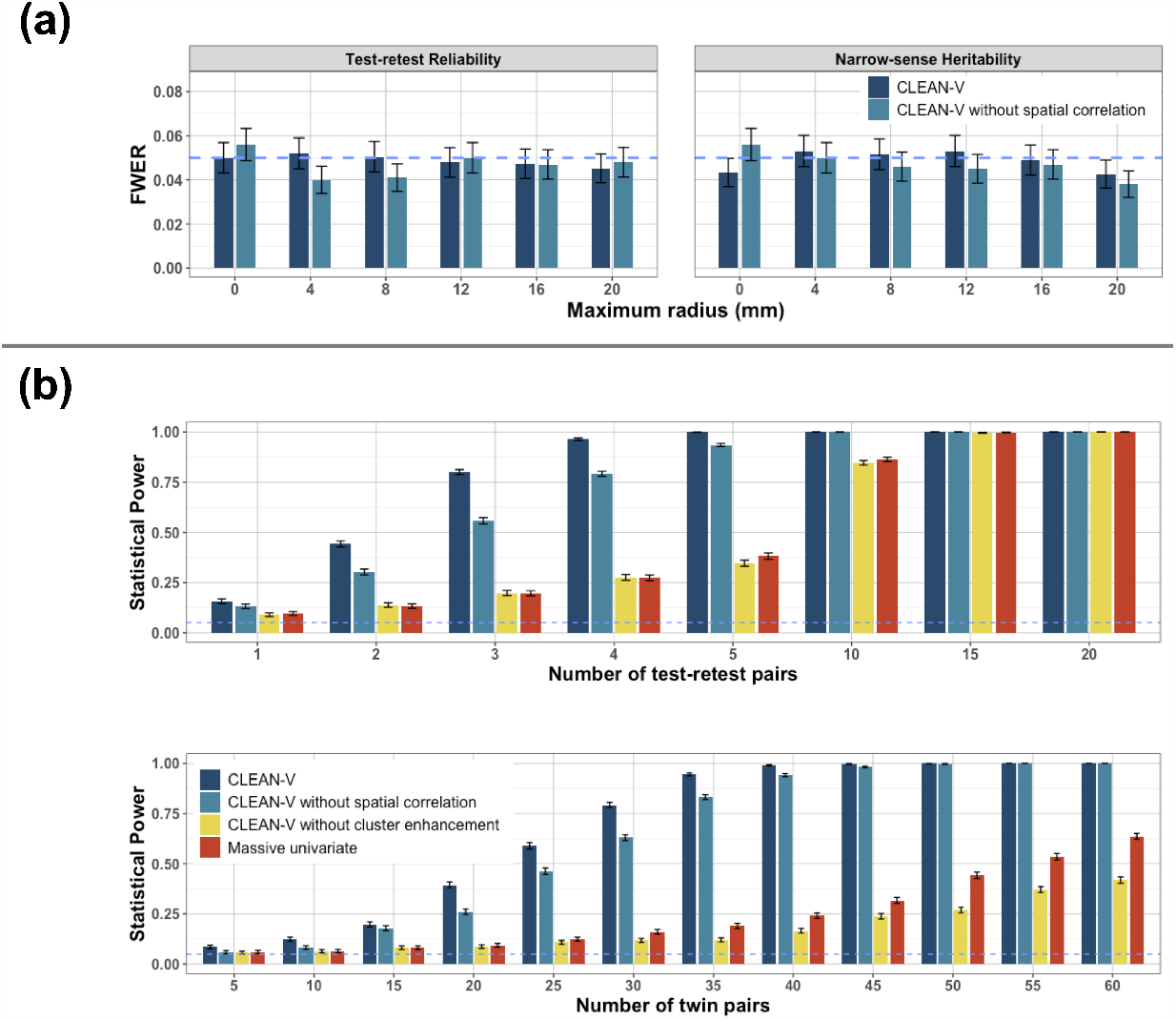
(a) Empirical FWER of CLEAN-V and its competitors under different maximum radii *r*_max_s for clusterwise enhancement from the data-driven simulation studies under the null hypothesis. Note that since massive univariate analysis does not involve the clusterwise procedure, massive univariate analysis is the same as CLEAN-V without spatial correlation with 0mm maximum radius. Similarly, CLEAN-V without cluster enhancement is equivalent to CLEAN-V with 0mm maximum radius. (b) Empirical power of CLEAN-V and its competitors under different signal-to-noise ratios. The error bars denote standard errors.

Figure 6(b) shows the power of the four competitors. The empirical power is calculated by the number of times rejecting the null hypothesis divided by the number of simulations (1000). We see that CLEAN-V’s power was uniformly higher than the competitors, and the power of CLEAN-V without spatial correlation was substantially higher than the massive univariate analysis. For test-retest reliability, however, the power of CLEAN-V without cluster enhancement showed a similar power as massive univariate analysis and even lower when the signal-to-noise ratio was high. Notably, CLEAN-V achieved strong power (80%) when the number of test-retest pairs was only 3. As the number of test-retest pairs increased to 5, almost 100% power was obtained by CLEAN-V. For narrow-sense reliability, CLEAN-V also achieved the highest power. Specifically, it reached 80% power when the number of twin pairs was 35. And it attained 100% power as the number of twin pairs increased to 40.

#### 3.5.2. Reproducibility

We computed vertex-level reproducibility from the simulation results. Specifically, for each vertex, we calculated the proportion of the vertex identified out of 1000 simulations. Figure 7(a) shows the reproducibility maps of three settings for both test-retest reliability and narrow-sense heritability. Clearly, CLEAN-V outperformed its competitors under all settings. CLEAN-V and CLEAN-V without spatial correlation identified substantial signal regions for test-retest reliability when sufficient test-retest pairs and twins were included. CLEAN-V was better than CLEAN-V without spatial correlation in all cases. Regardless of the number of test-retest pairs and twin pairs included in the samples of simulations, Massive univariate and CLEAN-V without cluster enhancement nearly cannot detect the signals of test-retest reliability and heritability. Because the signal region of heritability for the emotional task is relatively small, both CLEAN-V and CLEAN-V without spatial correlation identified small signal regions. However, CLEAN-V still identified more signal regions with higher reproducibility.

**Figure 7:**
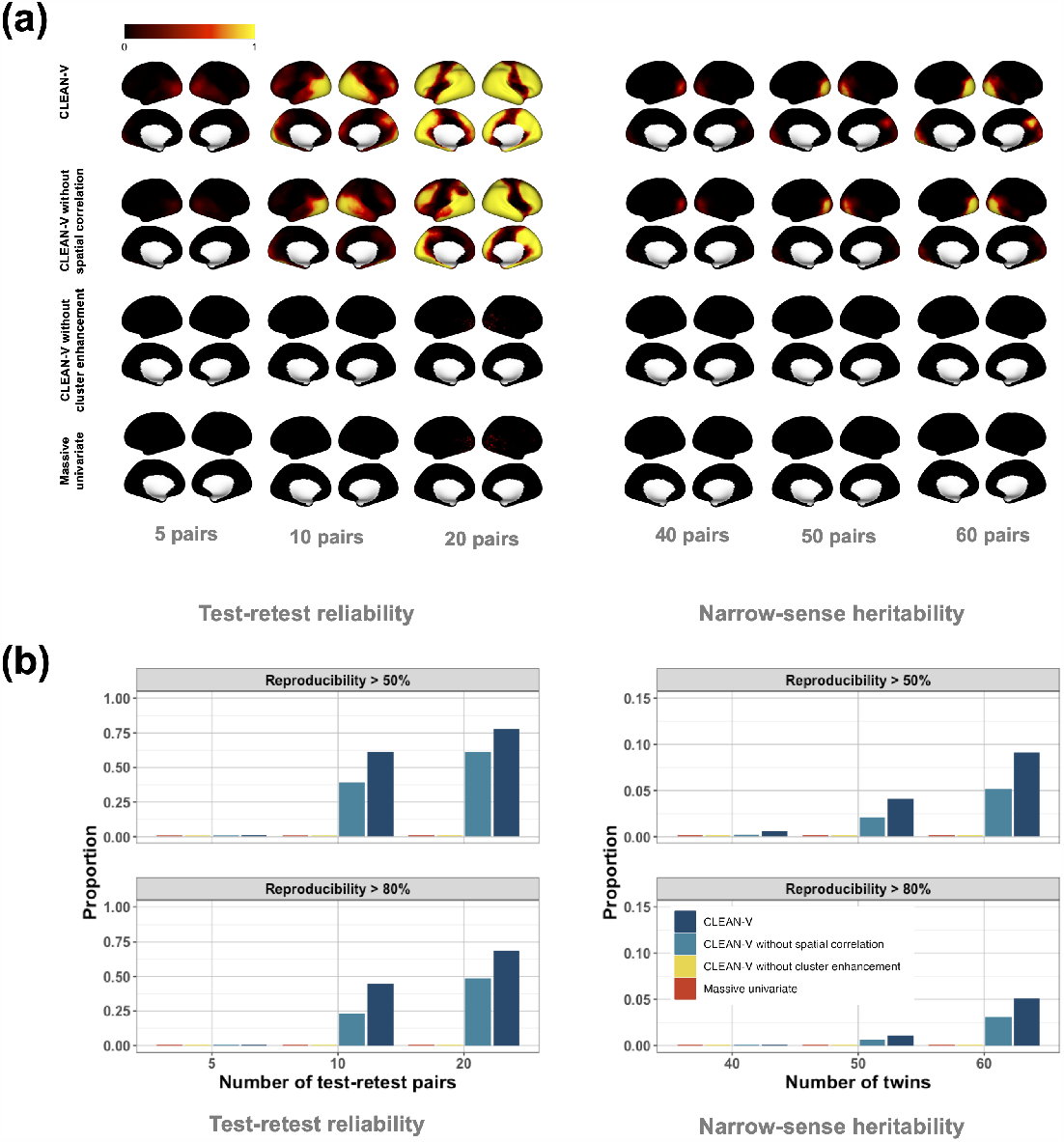
Reproducibility analysis from the data-driven simulation studies. The left and right figures describe the results for test-retest reliability and narrow-sense heritability. (a) The value of each vertex is the proportion of rejecting the null hypothesis among the 1000 simulated datasets. (b) Proportions of vertices with over 80% and 50% reproducibility for each method.

Figure 7(b) included the proportion of vertices whose reproducibility is over 80% or 50% for each approach. CLEAN-V outperformed in all scenarios. When there were 20 test-retest pairs in the sample for test-retest reliability analysis (all test-retest data), CLEAN-V remarkably produced 68% and 77% vertices over 80% and 50% reproducibility correspondingly, which were better than CLEAN-V without spatial correlation (48% and 61%) and considerably higher than CLEAN-V without cluster enhancement and Massive univariate (0% and 0.1% for both). For narrow-sense heritability analysis with 60 pairs of twins, 9% and 5% vertices had over 50% and 80% reproducibility from CLEAN-V, which is also notably higher than the proportion of vertices with CLEAN-V without spatial correlation (5% and 3%) and the other two methods (0% for both). For other settings, CLEAN-V also significantly outperformed the others.

## 4. Discussion

In this paper, we proposed a new method, CLEAN-V, for statistical inference and localization of test-retest reliability and narrow-sense heritability for task-based fMRI. Our method directly adjusts the global spatial autocorrelation, constructs spatially-enhanced test statistics in a locally powerful way, and controls FWER efficiently by permutations. We showed that CLEAN-V controlled familywise error rate and achieved high power simultaneously.

Our results showed that it is necessary to use both cluster-wise enhancement and spatial autocorrelation modeling to increase sensitivity in testing variance components, which aligns with previous findings [14, 15]. It was supported by comparing CLEAN-V to simpler methods: massive univariate analysis, CLEAN-V without spatial correlation, and CLEAN-V without cluster enhancement. The comparison results showed that there is no power improvement if only spatial dependence is considered (without cluster enhancement). Our cluster-wise enhancement method with univariate statistics led to increased statistical power by flexibly collecting neighbors’ information and adaptation. With a combination of the two modeling strategies (clusterwise enhancement and spatial autocorrelation modeling), CLEAN-V sufficiently detected test-retest reliability even if the proportion of test-retest data in samples was very small (15%) and achieved almost perfect power when the proportion increased to 25%. Considering that the number of images used in studies for test-retest reliability or heritability is mostly small (*N <* 100), CLEAN-V provides a powerful and sensitive approach that partially relaxes such concerns.

In the analysis of 5 experimental tasks of the HCP task-fMRI data, CLEAN-V, in general, distinguished the strength of test-retest reliability and narrow-sense heritability on the brain surface and among tasks. We also found a strong relationship between test-retest reliability and activation by comparing the CLEAN-V’s test-retest maps to activation maps. Thus, CLEAN-V offers possibilities to discover the correspondence between reliability regions and activation regions. Combining the score-based variance component test and permutations, CLEAN-V is much more computationally efficient than the classical likelihood-based tests since constructing the test statistics and finding FWER-controlled threshold only need fitting the null model once instead of refitting numerous models. The computational efficiency of CLEAN-V speaks of its practical utility.

This paper focused primarily on analyzing task-fMRI data. Still, CLEAN-V is widely applicable to many neuroimaging data types (e.g. cortical thickness) where ‘borrowing’ information across spatial domains is expected to improve power. Also, a comprehensive exploratory analysis by Weinstein et al. [15] reveals that applying exponential spatial autocorrelation function (SACF) is reasonable in many imaging modalities mapped onto the cortical surface. While applying the same assumption to volumetric (3D) imaging data could be questioned, we showed that a simplified version of CLEAN (‘CLEAN-V without spatial correlation’) also results in a dramatic increase in power and sensitivity compared to massive univariate analysis, which can be adopted easily.

Lastly, CLEAN-V provides useful insight into recent concerns on low test-retest reliability or heritability in neuroimaging studies. Although CLEAN-V does not provide effect size estimates, the fact that the adoption of ‘spatial variations’ resulted in increased power suggests that the current definition of test-retest reliability or heritability in neuroimaging studies might be too simple to characterize all sources of variations, such as spatial variations, which would provide limited understanding of variance components and lead to suboptimal estimations. For example, Risk and Zhu [6] showed how spatial modeling of heritability provides better estimates than mass univariate analysis. Extending the ‘inference’ problem of CLEAN-V to the ‘estimation’ problem is worth further investigations, which we leave as future work.

## 5. Software

Without parallel computing, implementing CLEAN-V for 342 images (each image containing 10,000 vertices) with 5,000 permutations took less than 10 minutes on a laptop, which supports the practical utility of the method. We are working on additional optimization of the software with parallel computing, which is publicly available at https://github.com/junjypark/CLEAN.

## Declaration of Competing Interest

None.

### Acknowledgement

We would like to thank reviewers for their helpful comments and suggestions. EWD receives funding from the Brain and Behavior Research Foundation Young Investigator Award (NARSAD), the Canadian Institutes of Health Research. CH receives research funding from the National Institute of Mental Health (NIMH), the Brain and Behavior Research Foundation, the University of Toronto, and the Centre for Addiction and Mental Health (CAMH) Foundation. NR receives funding from the Natural Sciences and Engineering Research Council of Canada (RGPIN-2020-05897). ANV receives funding from the National Institute of Mental Health (1/3R01MH102324 & 1/5R01MH114970), Canadian Institutes of Health Research, Canada Foundation for Innovation, CAMH Foundation, and University of Toronto. JYP receives funding from the Natural Sciences and Engineering Research Council of Canada (RGPIN-2022-04831), and the University of Toronto’s Data Science Institute, McLaughlin Centre, and Connaught Fund.

Data were provided by the Human Connectome Project, WU-Minn Consortium (Principal Investigators: David Van Essen and Kamil Ugurbil; 1U54MH091657) funded by the 16 NIH Institutes and Centers that support the NIH Blueprint for Neuroscience Research; and by the McDonnell Center for Systems Neuroscience at Washington University. This research was supported in part by

NIH grants R01EB016061 and R01EB026549 from the National Institute of Biomedical Imaging and Bioengineering and R01 MH116026 from the National Institute of Mental Health.

## Supplementary materials

## A. Normal approximation of mixture chi-square distribution

Combing A.1 and A.2, the null distribution of 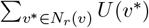is approximated by the normal distribution when *N* is large. This implies that *T*_*r*_(*v*) (the standardized 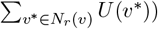 follows the standard normal distribution approximately under *H*_0_.

### A.1 Proof that 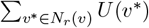 *follows mixture chi-square under H*_0_

Based on our model specification, we define **y** = (*y*_1_(1), *y*_2_(1), …, *y*_*N*_ (1), …, *y*_1_(*V*), *y*_2_(*V*), …, *y*_*N*_ (*V*))′, which leads to

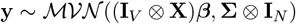

where *β* = (*β*(1)′, …, *β*(*V*)′)′ and ⊗ denotes for the Kronecker product. Because OLS is used to estimate *β*(*v*), we have

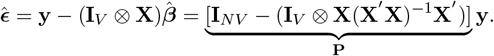

After estimating 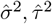 and 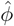 by covariance regression analysis, we get 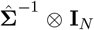 where 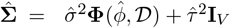. Then, we can get the 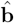 by estimated conditional expectation 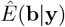from the joint distribution of (b, y):

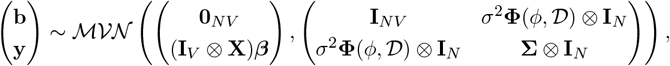

which provides

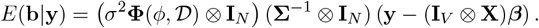

Plugging in the estimated spatial parameters, we obtain

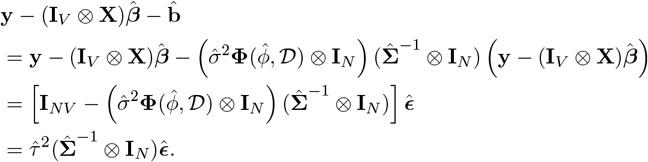

To get corresponding test statistics for each vertice *v*, we define **S**_*v*_ : ℝ*NV* → ℝ*N* to identify the subset of all images’ element for vertice 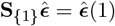. Then, the test statistic becomes

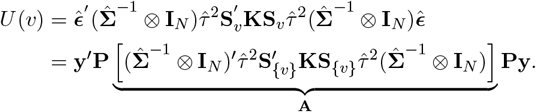

Similarly, define 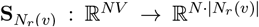 to identify the subset of all subjects’ elements for vertices in 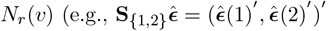. Then, for cluster-enhanced test statistic, we have

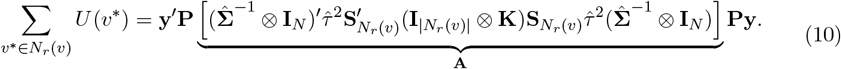

Since **Py** follows multivariate normal with zero mean vector and the covariance matrix **P**(**Σ**⊗**I**_*N*_)**P, y**′ **PAPy** follows a mixture chi-square distribution with weights being the eigenvalues of **QAQ**′ where **Q** corresponds to the Cholesky decomposition of 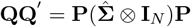 [31].

### A.2 Proof that the mixture chi-square distribution converges in distribution to a normal distribution

We use the Lyapunov Central Limit Theorem to prove the mixture chi-square distribution converges in distribution to a normal distribution as the number of components in the mixture chi-square goes to infinity. Assume {*Y*_1_, …, *Y*_*k*_} is a sequence of independent random variables, each with finite expected value μi and variance 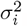, and 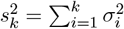. The Lyapunov Central Limit Theorem states that if the following condition is satisfied for the sequence:

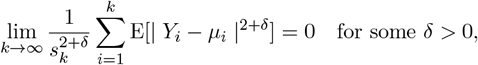

then 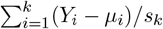 will converge weakly to the standard normal distribution.

Suppose 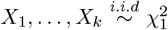 are the components of the mixture chi-square for our test statistic (10) with their corresponding weights *λ*_1_, …, *λ*_*k*_ which are the eigenvalues of **QAQ**′. Then it is sufficient to prove that the sequence of *λ*_*i*_*X*_*i*_ satisfies the Lyapunov’s condition. Because the fourth central moment of 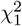 is 48, we can get the fourth central moment of *λ*_*i*_*X*_*i*_: 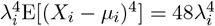. Then, choosing *δ* = 2, we get

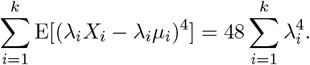

Because Var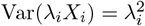Var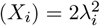, we have

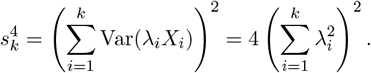

The result above leads to

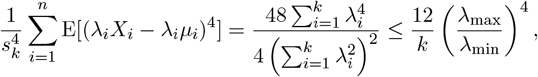

where *λ*_max_ and *λ*_min_ are the largest and smallest eigenvalues, respectively. As *k* goes to infinity,

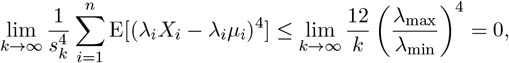

provided that *λ*_max_*/λ*_min_ is bounded. Therefore, the Lyapunov’s condition is satisfied and 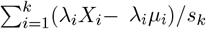converges in distribution to a standard normal random variable which is equivalent to 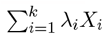 converges weakly to a normal distribution.

## B. Proof that *T*_*r*_(*v*) is constructed by assuming *θ*^*2*^(*v*^*∗*^)s in the neighborhood *N*_*r*_(*v*) are the same

Let *v*_1_, …, *v*|*N*_*r*_ (*v*)| be indices of vertices in *N*_*r*_(*v*). From our model assumptions, the model within *N*_*r*_(*v*) is

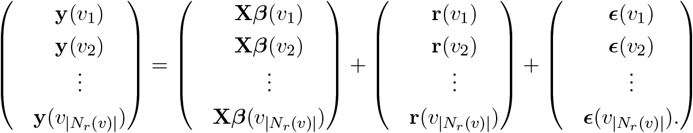

Assuming that *θ*^2^(*v*^∗^)s in *N*_*r*_(*v*) have the same value *θ*^2^, then

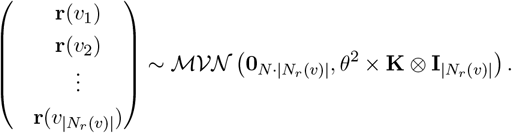

Since the 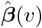s and 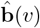are obtained from the null model, they keep the same estimated value as we described in section 2.3. Within the *N*_*r*_(*v*), the score-based variance component test statistic for *θ*^2^ becomes

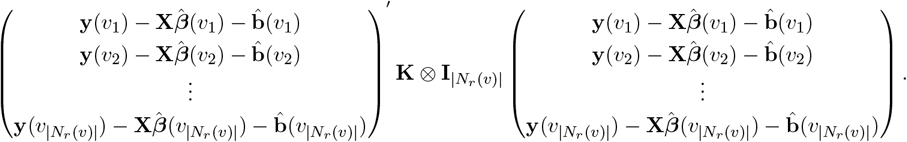

Note that, this matrix multiplication is summarized by

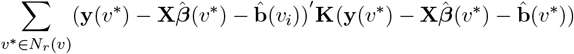

which is equal to 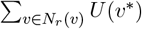. This result suggests that *T*_*r*_(*v*) is constructed by assuming the *θ*^2^(*v*)*s* are same in the neighborhood *N*_*r*_(*v*) as we stated in Section 2.3.

## C. Supplementary figures

Figures S1 and S2 show binarized maps of statistical significance when CLEAN-V, CLEAN-V without spatial correlation, CLEAN-V without cluster enhancement, and massive univariate analysis are applied to language, social cognition, relational processing, and gambling tasks.

**Figure S1:**
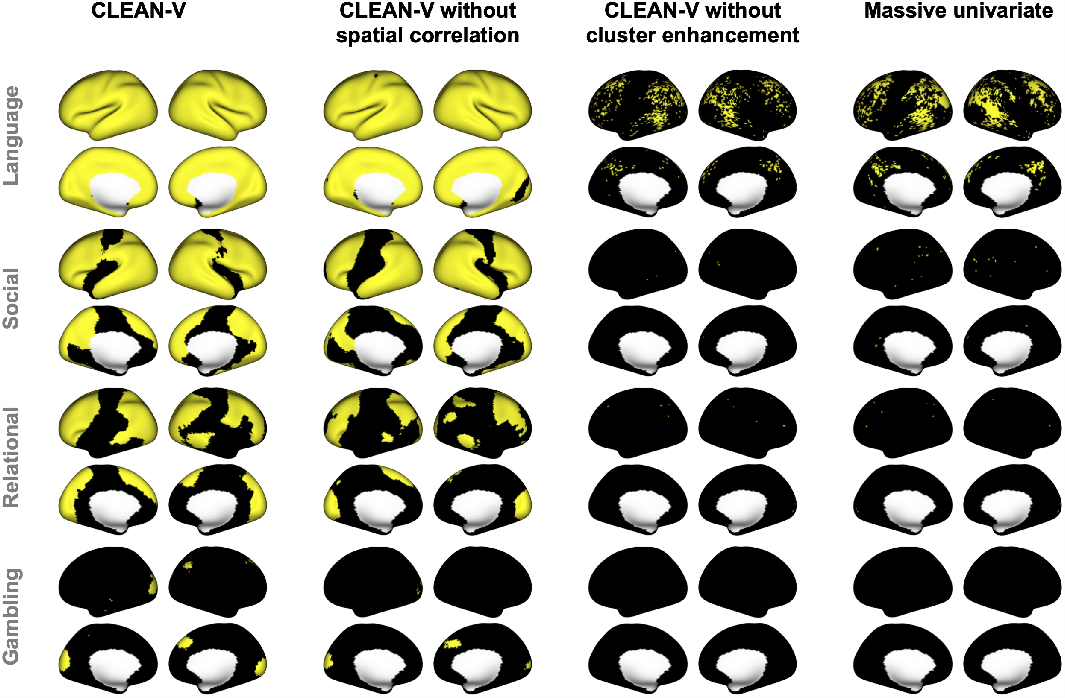
Test-retest reliability localization results from the four methods for the other four tasks.

**Figure S2:**
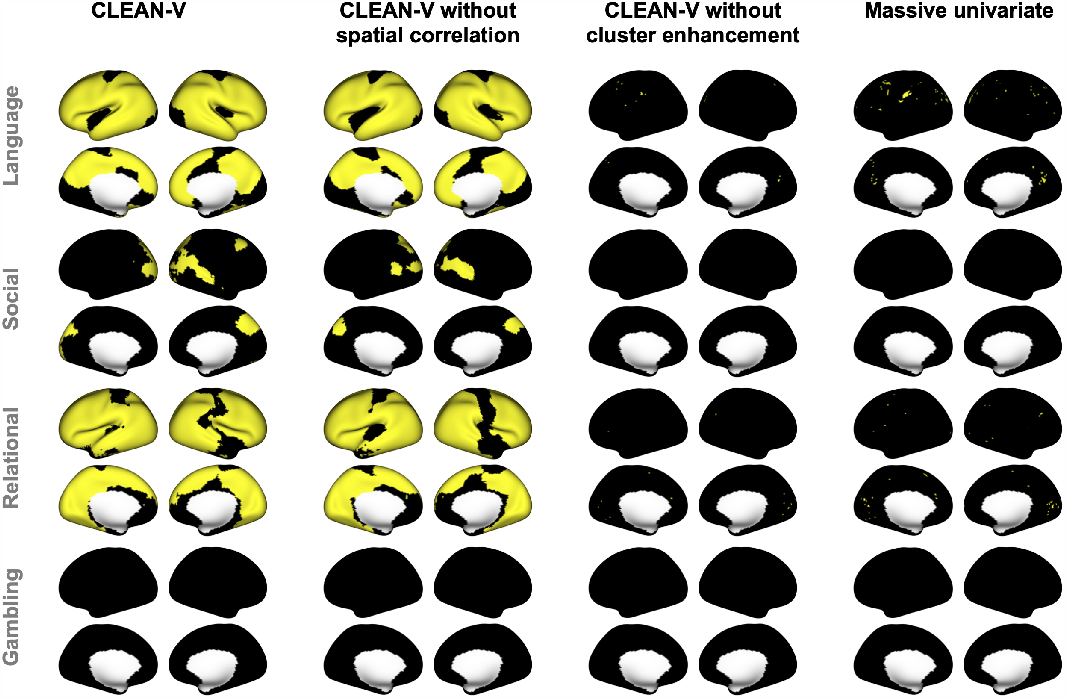
Narrow-sense heritability localization results from the four methods for the four other tasks.

## References

[1] M. L. Elliott, A. R. Knodt, D. Ireland, M. L. Morris, R. Poulton, S. Ramrakha, M. L. Sison, T. E. Moffitt, A. Caspi, A. R. Hariri, What is the test-retest reliability of common task-functional MRI measures? New empirical evidence and a meta-analysis, Psychological Science 31 (2020) 792–806.

[2] J. T. Kennedy, M. P. Harms, O. Korucuoglu, S. V. Astafiev, D. M. Barch, W. K. Thompson, J. M. Bjork, A. P. Anokhin, Reliability and stability challenges in ABCD task fMRI data, NeuroImage 252 (2022) 119046.

[3] S. Marek, B. Tervo-Clemmens, F. J. Calabro, D. F. Montez, B. P. Kay, A. S. Hatoum, M. R. Donohue, W. Foran, R. L. Miller, T. J. Hendrickson, et al., Reproducible brain-wide association studies require thousands of individuals, Nature 603 (2022) 654–660.

[4] G. A. Blokland, K. L. McMahon, P. M. Thompson, N. G. Martin, G. I. de Zubicaray, M. J. Wright, Heritability of working memory brain activation, Journal of Neuroscience 31 (2011) 10882–10890.

[5] M. A. Lindquist, J. Spicer, I. Asllani, T. D. Wager, Estimating and testing variance components in a multi-level GLM, NeuroImage 59 (2012) 490–501.

[6] B. B. Risk, H. Zhu, ACE of space: estimating genetic components of high-dimensional imaging data, Biostatistics 22 (2021) 131–147.

[7] S. Noble, D. Scheinost, R. T. Constable, A guide to the measurement and interpretation of fMRI test-retest reliability, Current Opinion in Behavioral Sciences 40 (2021) 27–32.

[8] S. M. Smith, T. E. Nichols, Threshold-free cluster enhancement: Addressing problems of smoothing, threshold dependence and localisation in cluster inference, NeuroImage 44 (2009) 83–98.

[9] J. Y. Park, M. Fiecas, A. D. N. Initiative, et al., Permutation-based inference for spatially localized signals in longitudinal MRI data, NeuroImage 239 (2021) 118312.

[10] F. Mejia, Y. Yue, D. Bolin, F. Lindgren, M. A. Lindquist, A Bayesian general linear modeling approach to cortical surface fMRI data analysis, Journal of the American Statistical Association 115 (2020) 501–520.

[11] D. Spencer, Y. R. Yue, D. Bolin, S. Ryan, A. F. Mejia, Spatial Bayesian GLM on the cortical surface produces reliable task activations in individuals and groups, NeuroImage 249 (2022) 118908.

[12] J. L. Bernal-Rusiel, M. Reuter, D. N. Greve, B. Fischl, M. R. Sabuncu, A. D. N. Initiative, et al., Spatiotemporal linear mixed effects modeling for the mass-univariate analysis of longitudinal neuroimage data, NeuroImage 81 (2013) 358–370.

[13] B. B. Risk, D. S. Matteson, R. N. Spreng, D. Ruppert, Spatiotemporal mixed modeling of multi-subject task fMRI via method of moments, NeuroImage 142 (2016) 280–292.

[14] J. Y. Park, M. Fiecas, CLEAN: Leveraging spatial autocorrelation in neuroimaging data in clusterwise inference, NeuroImage 255 (2022) 119192.

[15] S. M. Weinstein, S. N. Vandekar, E. B. Baller, D. Tu, A. Adebimpe, T. M. Tapera, R. C. Gur, R. E. Gur, J. A. Detre, A. Raznahan, et al., Spatially-enhanced clusterwise inference for testing and localizing intermodal correspondence, NeuroImage (2022) 119712.

[16] M. A. Lindquist, The statistical analysis of fMRI data, Statistical Science 23 (2008) 439 –464.

[17] M. M. Monti, Statistical analysis of fMRI time-series: a critical review of the GLM approach, Frontiers in human neuroscience 5 (2011) 28.

[18] H. Ganjgahi, A. M. Winkler, D. C. Glahn, J. Blangero, P. Kochunov, T. E. Nichols, Fast and powerful heritability inference for family-based neuroimaging studies, NeuroImage 115 (2015) 256–268.

[19] T. Ge, T. E. Nichols, P. H. Lee, A. J. Holmes, J. L. Roffman, R. L. Buckner, M. R. Sabuncu, J. W. Smoller, Massively expedited genome-wide heritability analysis (MEGHA), Proceedings of the National Academy of Sciences 112 (2015) 2479–2484.

[20] M. C. Wu, S. Lee, T. Cai, Y. Li, M. Boehnke, X. Lin, Rare-variant association testing for sequencing data with the sequence kernel association test, The American Journal of Human Genetics 89 (2011) 82–93.

[21] T. Zou, W. Lan, H. Wang, C.-L. Tsai, Covariance regression analysis, Journal of the American Statistical Association 112 (2017) 266–281.

[22] Z. Li, Y. Liu, X. Lin, Simultaneous detection of signal regions using quadratic scan statistics with applications to whole genome association studies, Journal of the American Statistical Association 117 (2022) 823–834.

[23] Datta, S. Banerjee, A. O. Finley, A. E. Gelfand, Hierarchical nearest-neighbor gaussian process models for large geostatistical datasets, Journal of the American Statistical Association 111 (2016) 800–812.

[24] D. M. Barch, G. C. Burgess, M. P. Harms, S. E. Petersen, B. L. Schlaggar, M. Corbetta, M. F. Glasser, S. Curtiss, S. Dixit, C. Feldt, et al., Function in the human connectome: task-fMRI and individual differences in behavior, NeuroImage 80 (2013) 169–189.

[25] M. F. Glasser, S. N. Sotiropoulos, J. A. Wilson, T. S. Coalson, B. Fischl, J. L. Andersson, J. Xu, S. Jbabdi, M. Webster, J. R. Polimeni, D. C. Van Essen, M. Jenkinson, The minimal preprocessing pipelines for the Human Connectome Project, NeuroImage 80 (2013) 105–124.

[26] D. D. Pham, J. Muschelli, A. F. Mejia, ciftitools: A package for reading, writing, visualizing, and manipulating cifti files in R, NeuroImage 250 (2022) 118877.

[27] C. Hawco, E. W. Dickie, G. Jacobs, Z. J. Daskalakis, A. N. Voineskos, Moving beyond the mean: Subgroups and dimensions of brain activity and cognitive performance across domains, NeuroImage 231 (2021) 117823.

[28] S. M. Weinstein, S. N. Vandekar, A. Adebimpe, T. M. Tapera, T. Robert-Fitzgerald, R. C. Gur, R. E. Gur, A. Raznahan, T. D. Satterthwaite, A. F. Alexander-Bloch, et al., A simple permutation-based test of intermodal correspondence, Human brain mapping 42 (2021) 5175–5187.

[29] S. Geuter, G. Qi, R. C. Welsh, T. D. Wager, M. A. Lindquist, Effect size and power in fMRI group analysis, Biorxiv (2018) 295048.

[30] Eklund, T. E. Nichols, H. Knutsson, Cluster failure: Why fMRI inferences for spatial extent have inflated false-positive rates, Proceedings of the national academy of sciences 113 (2016) 7900–7905.

[31] P. Duchesne, P. L. De Micheaux, Computing the distribution of quadratic forms: Further comparisons between the liu–tang–zhang approximation and exact methods, Computational Statistics & Data Analysis 54 (2010) 858–862.

